# The Turbulent Brain: a toolbox to compute the turbulent dynamical behaviour of whole-brain activity

**DOI:** 10.1101/2025.11.14.688068

**Authors:** Yonatan Sanz Perl, Jakub Vohryzek, Anira Escrichs, Morten L. Kringelbach, Gustavo Deco

**Affiliations:** Center for Brain and Cognition, Computational Neuroscience Group, Universitat Pompeu Fabra, Barcelona, Spain; National Scientific and Technical Research Council (CONICET), CABA, Buenos Aires, Argentina; Department of Engineering, Universidad de San Andrés, Buenos Aires, Argentina; International Centre for Flourishing, Universities of Oxford (UK), Aarhus (Denmark) and Pompeu Fabra (Spain); Department of Psychology & Centre for Cognitive Neuroscience, Paris-Lodron-University Salzburg, Salzburg, Austria; Department of Psychiatry, University of Oxford, Oxford, United Kingdom; Center for Music in the Brain, Department of Clinical Medicine, Aarhus University, Århus, Denmark; Life and Health Sciences Research Institute (ICVS), School of Medicine, University of Minho, Braga, Portugal; Centre for Eudaimonia and Human Flourishing, University of Oxford, Oxford, United Kingdom; Department of Information and Communication Technologies, Universitat Pompeu Fabra, Barcelona, Spain; Institució Catalana de la Recerca i Estudis Avancats (ICREA), Barcelona, Spain

**Author notes:** These authors contribute equally (Yonatan Sanz Perl, Jakub Vohryzek).

## Abstract

Brain disorders remain one of the greatest challenges in neuroscience. Computational precision medicine offers a promising framework to reveal the mechanistic principles governing both healthy and pathological brain dynamics. In this context, we present The Turbulent Brain (TTB) toolbox, an open-source MATLAB platform that implements and disseminates the turbulence framework for immediate and widespread application in neuroscience research. The toolbox combines model-free and model-based analyses through an intuitive graphical interface, enabling customizable workflows across different parcellations and preprocessing pipelines. Our toolbox builds on recent advances demonstrating that turbulence physics can effectively capture the complex spatiotemporal dynamics of brain activity as a high-dimensional system operating far from equilibrium. This framework allows researchers to explore how fundamental mechanisms are modulated across distinct brain states and neurological and psychiatric disorders, including disorders of consciousness and major depressive disorder. As an illustrative application case, we used TTB to analyse fMRI data from 176 individuals from the Human Connectome Project during resting state and movie watching, showing that movie watching, as compared to resting state, significantly reduces information transmission in terms of turbulence measures, with stronger reductions observed during Creative Commons movie watching compared to Hollywood movies. In summary, the TTB toolbox establishes an accessible framework providing a methodological bridge between nonlinear physics and computational neuroscience, offering new tools for investigating psychiatric conditions and altered states of consciousness.

## Introduction

Brain activity, even at rest, exhibits rich spatio-temporal dynamics. Recent research has demonstrated that the physics of turbulence can capture whole-brain activity^1,2^. This parallelism is important as it offers a principled framework for describing the brain as a high-dimensional dynamical system operating far from equilibrium. Turbulence allows us to investigate the mechanism underlying the brain’s ability for optimal and time-critical information processing as well as fast reactiveness to external stimuli. At the large scale, rather than focusing solely on localized activity or statistical dependence between brain regions, known as functional connectivity, the turbulent regime captures how macroscopic neural interactions, in terms of emerging and dissolving transient synchronization structures, evolve over time and space supported by the brain’s white-matter connectivity - the connectome^3^.

Turbulence was first documented by Leonardo da Vinci and has been studied since as a fundamental phenomenon in fluid dynamics^4^. It attempts to describe the whirls observed in river flows as complex, multiscale spatiotemporal patterns that arise when a system is driven far from equilibrium. In turbulent flows, these patterns emerge as energy is transferred from large scales down to smaller scales, leading to rapid and optimal mixing, broadband spectral content, and scale-free statistical properties. To capture this cascading process, Kolmogorov developed the phenomenological theory of turbulence which introduced the concept of the structure function to describe how correlations between any two points in a system decay with spatial distance revealing power-law scaling in the inertial subrange of the system’s spectrum **(Figure 1A)**^5,6^. Beyond the field of fluid dynamics, Kuramoto’s work has generalised the concept of turbulence to other physical systems and in particular coupled oscillators. By measuring the local synchronization across space and time in a network of coupled-oscillators, Kuramoto demonstrated the emergence of turbulent-like complex spatio-temporal patterns. Akin to vortices in fluids, these local synchrony fluctuations capture how information is mixed and redistributed throughout the network of coupled-oscillators^7,8^. These spiral-like vortices in fluid dynamics and systems of coupled oscillators have also been demonstrated in brain activity, where oscillatory vortices form around their phase singularity centers^9,10^. Importantly, these can be quantified using the same formalism of local synchronization in terms of the Kuramoto Order Parameter ^2^ **(Figure 1B)**.

**Figure 1:**
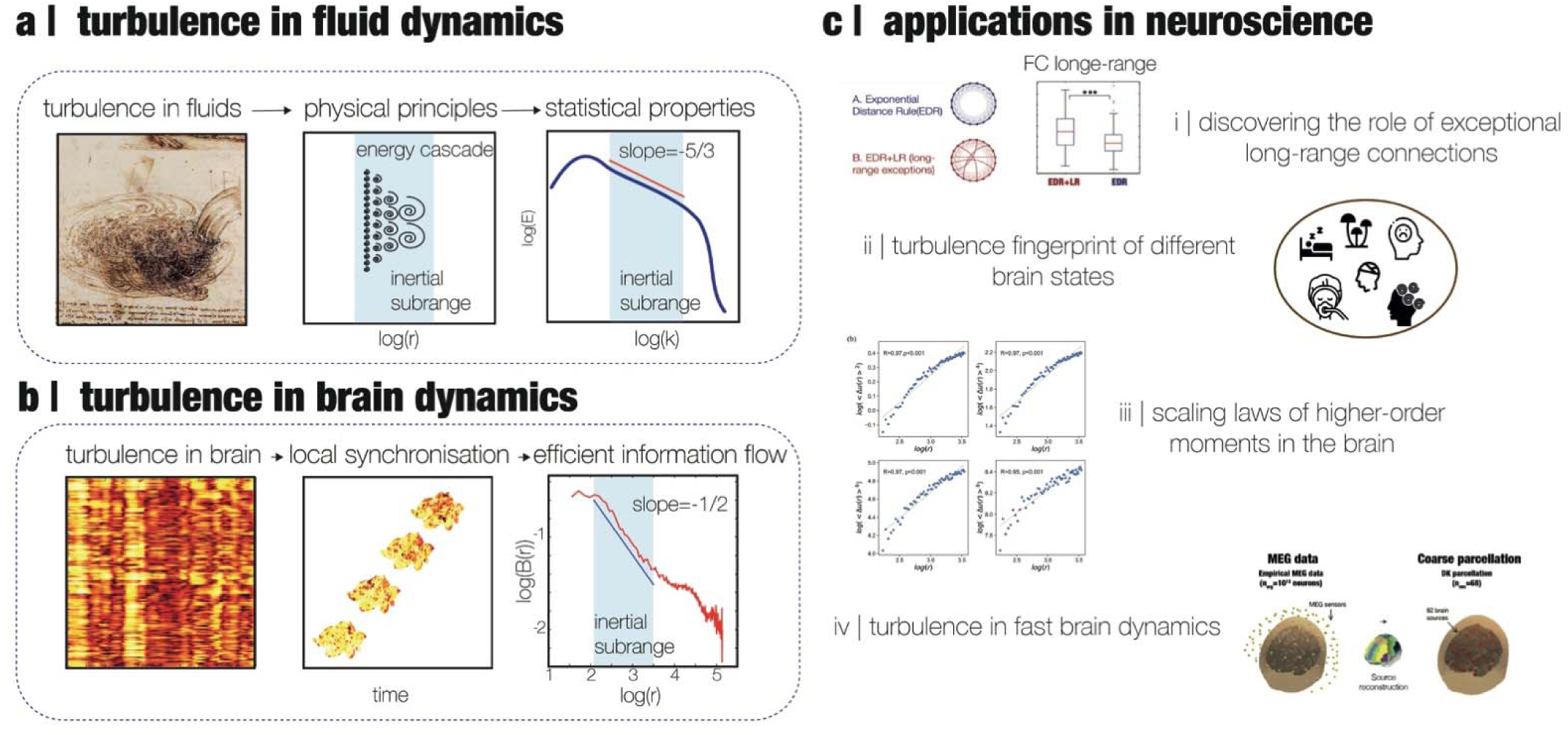
***Turbulence in brain and neuroscience applications. a| Turbulence in fluids:*** *The study of turbulence in fluid dynamics inspired Leonardo da Vinci to coin the term “turbolenza” for the whirls of chaotic movement that promoteoptimal mixing across space and time. **Physical principles:** Kolmogorov’s phenomenological theory of turbulence introduced the concept of structure function, inspired by Richardson’s idea of cascading eddies, to describe the statistical properties of high-dimensional non-linear fluid dynamics. **Statistical properties:** Such formalism allowed for the discovery of power laws in the inertial subrange with structure function showing a universal energy scaling of k^-^*^5^*^/^*^3^ *where k is the wave number of the spectral scale, demonstrating how the energy/information transfer cascade, found in turbulence, takes place. **b| Turbulence in brain:** Recently, work by Kuramoto used coupled-oscillators to show turbulence in the whirls of synchronised oscillators promoting optimal mixing properties. **Local synchronisation:** The turbulence in non-fluid dynamics was derived from the local Kuramoto order parameter in terms of the level of amplitude turbulence. In the brain, such a type of turbulence can be derived from local phases of the analytical signal (defined for several brain regions at a specific spatial kernel). **Efficient information flow:** At the inertial subrange, it has been shown to follow a power-law for the correlation B(r) as a function of **r** reflecting the optimality of efficiency transfer in the spacetime information flow in brain dynamics. **c|** The versatility of the turbulence framework has allowed for several complementary applications to neuroscience. **i) Structural backbone supporting turbulent brain dynamics:** Recent work has demonstrated the importance of long-range exceptions beyond and above the exponential distance rule (EDR) of connectivity wiring for optimal turbulent properties of spatio-temporal brain dynamics. **ii) Turbulent signatures of different brain states.** Changes to turbulent-like brain dynamics have been observed in different brain states such as sleep, meditation, anaesthesia, disorders of consciousness and psychedelics. **iii) Scaling laws of higher-order moments in the brain** Turbulent-like brain activity exhibits scaling in higher-order moments of the local Kuramoto order parameter important for the brain’s information transmission and reactivity. **iv) MEG-based signatures of turbulence-like brain dynamics:** Turbulent-like dynamics have been also observed across temporal scale in MEG*.

The turbulence framework has been remarkably versatile^11^ finding its use in a wide range of neuroscientific applications (**Figure 1C**). At the microscopic level Sheremet and colleagues have found turbulence and energy cascades in the hippocampus^12^. At the macroscopic level, the interaction of these turbulent vortices expresses the levels of local synchronisation in the underlying brain signals and it has been used to describe the information processing underlying the necessary computation for developing cognition^13^. This vortex space offers an excellent abstraction for capturing the mechanisms underlying the orchestration of brain function over spacetime. In this context, recent studies on the structure-function relationship of brain organisation^14^, have shown that long-range anatomical connections that deviate from the known exponential distance rule (EDR)^15^ play a critical role in supporting optimal turbulent properties of large-scale brain dynamics ^10^. These rare connections act as shortcuts for information flow, enhancing the efficiency of multiscale communication ^16^. Moreover, using measures derived from the local Kuramoto Order Parameter (KOP), researchers have demonstrated systematic differences in turbulence levels across various brain states such as sleep, anaesthesia, disorders of consciousness and psychedelic states. For example, conscious wakefulness is typically associated with optimal levels of spatiotemporal variability in local synchrony - interpreted as a hallmark of rich, metastable dynamics—whereas unconscious states often show a breakdown of this turbulent structure^17^. An opposite trend is observed under the influence of classical serotonergic psychedelics where the level of turbulence is increased^18^. Turbulent-like brain activity also exhibits scaling behaviour in higher-order statistical moments of the local KOP allowing for a better description of information transmission and reactivity of the brain^19^. Lastly, these turbulent-like dynamics are not confined to fMRI but have also been observed in MEG data, where they span a broad range of temporal scales^20^. In a recent work, we exhaustively reviewed the applications of the turbulent brain approach for different neuroscientific questions^3^.

To quantify the turbulence in neuroimaging data we have implemented two complementary approaches developed in previous works, the model-free and model-based framework.

In the model-free framework, we use the local KOP to quantify turbulence as the variability of local synchronization across space and time^7,21^, analogous to rotational vortices in fluid dynamics whose sizes define different information-processing scales. It follows that we operate in ‘vortex space’ rather than signal space to assess information transfer in the brain ^22^. Practically, the spatial and temporal scales of the turbulence are defined by the local kernel based on the exponential distance rule and, and by the temporal acquisition of the data leveraging the instantaneous phase relationship between brain regions, respectively. From this spatially and temporally rich ‘vortex space’ several measures of turbulence can be derived. At the global level, turbulence is measured as the variability of the local KOP across space and time, and the transfer correlation quantifies the correlation of local synchronisation across space at different scales. The turbulence regime also endows the brain with an efficient information cascade measured as the correlation of the local level of synchronisation across scales (Information Cascade Flow). The average across scales of the information cascade flow is defined as the Information cascade. Furthermore, given the multiscale nature of turbulence in brain dynamics, these measures can be computed at the resting-state network and regional level **(Figure 2A)**.

**Figure 2:**
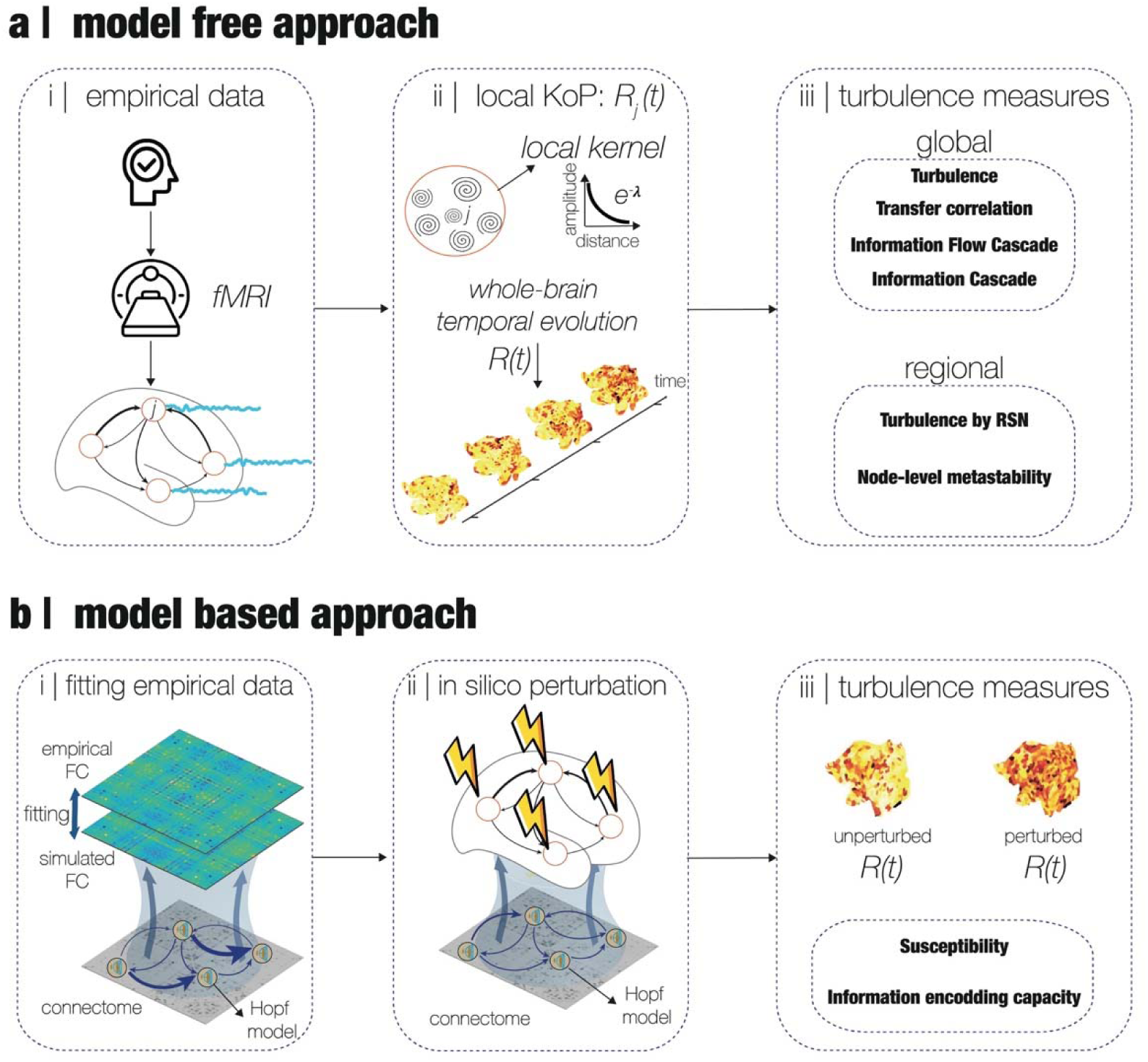
***Quantifying turbulence in global brain states using model-free and model-based frameworks.*** *Despite the microscopic complexity of brain activity, global brain states such as wakefulness, sleep, anaesthesia, and altered states of consciousness can be characterized by their macroscopic spatiotemporal dynamics. To quantify these dynamics, we implemented two complementary approaches. **a| Model-free approach:** Starting from empirical fMRI data (i), we computed the local Kuramoto Order Parameter (KOP) R_j_(t). The local KOP is using a spatially decaying kernel (ii), capturing the whole-brain temporal evolution of local synchronisation. This allows estimation of global and regional turbulence metrics (iii), including turbulence, transfer correlation, information cascades, and RSN-level metastability. **b| Model-based approach:** (i) We fitted whole-brain Hopf models to the empirical functional connectivity by adjusting the global coupling parameter G and then subjected the fitted model to in silico perturbations (ii). This enabled the assessment of turbulence-based sensitivity metrics (iii), including susceptibility and information encoding capacity, which reflect the system’s responsiveness to external stimuli and its capacity for information processing. Together, this framework reveals how turbulent dynamics support and distinguish global brain states*.

For the model-based framework, we used whole-brain modelling with coupled non-linear oscillators that integrates the anatomy and local dynamics of brain activity^23–25^. Specifically, we used the well-known Hopf whole-brain model where each region is described by a non-linear Stuart-Landau oscillator ^26,27^. The model has a few free parameters, the local bifurcation parameter that determines the dynamical regimes of the local dynamics, and the global coupling parameter which globally scales the weights of the structural connectivity representing the number of fibers connecting brain regions. By setting the bifurcation parameter at the edge of the Hopf bifurcation^28,29^, while optimizing the global coupling parameter, the model adequately reflects many properties of rich spontaneous brain dynamics including turbulence. Importantly, in-silico perturbations on the fitted whole-brain model allow the computation of two key measures: information encoding capability, which is maximal when the brain operates in turbulent regime ^2^, and susceptibility which is enhanced by rare long-range anatomical connections in the brain ^10^ **(Figure 2B)**.

Collectively, these results align the turbulence framework with the growing focus on computational precision medicine as a promising avenue for uncovering the mechanistic principles governing both healthy and pathological brain dynamics^30–32^. Here, we present The Turbulent Brain Toolbox (TTB), an open-source Matlab toolbox that implements an easily computable pipeline for the model-free and model-based analyses built upon the turbulent brain dynamics framework ^3^. Our goal is to further foster the implementation of the turbulent brain analysis by releasing a simple toolbox with a graphical user interface offering additional optional computations and allowing the user to customise their own analysis. While the mathematical foundations of turbulent brain dynamics analysis are not straightforward, this toolbox is designed to make their application accessible to a broad range of practitioners. Furthermore, the accompanying step-by-step tutorial is intended to guide the practitioners through the process, enabling them to swiftly gain proficiency in the derivation and comparison of turbulent measures across various brain states. This practical approach aims to lower the barrier to adoption and encourages a wider use of turbulence-based analyses in neuroscience, while remaining adaptable to diverse parcellations and preprocessing pipelines.

## Results

### The Turbulent Brain Toolbox (TTB)

The Turbulent Brain (TTB) is an open-science MATLAB toolbox that allows users to compute both model-free and model-based analyses based on the turbulence brain hypothesis. The toolbox is freely available at: https://github.com/yonisanzperl/The-Turbulent-Brain-v2. We implemented TTB as a pipeline in MATLAB version 2024a (The MathWorks, Natick, USA), and it does not require any additional packages except for the Statistical Parametric Mapping (SPM) toolbox (https://www.fil.ion.ucl.ac.uk/spm/), which is used to generate some of the graphical outputs. The toolbox can be installed in the user’s preferred directory, which also serves as the working folder where the results will be stored. Input data, however, can be located anywhere on the user’s system. TTB includes a graphical user interface (GUI) to guide users through the different steps of the framework, allowing them to perform model-free and model-based analyses independently. The toolbox is designed for neuroscientists and clinical researchers, enabling exploration of turbulence-related measures of brain dynamics without requiring a background in computational modeling or mathematics.

The user input requirements for the basic workflows of the toolbox are depicted in **Figure 3a**. The upper row describes the inputs needed for the model-free approach: i) the functional data (fMRI) for each brain condition and participant, using a fine-grained brain parcellation (such as Schaefer 1000^33^); ii) the center of mass (CoG) of each brain region in the selected parcellation; iii) the assignment of each brain region to functional resting-state networks (RSNs) in the parcellation (e.g., Yeo 7 RSN ^34^); iv) a set of parameters, which can be divided into fMRI parameters (e.g., TR and timepoints), filtering parameters, and spatial parameters. In the lower row of **Figure 3a**, the inputs for the model-based approach are shown. These are essentially the same as those required for the model-free approach, with two additions: a group-average structural connectome aligned to the same parcellation as the fMRI data, used for constructing whole-brain models, and model parameters, such as the range of values to explore for the coupling strength parameter (**G**). The toolbox includes a template database containing the CoG, group-average structural connectivity from healthy participants from the Human Connectome Project (see Methods), and RSN assignments for the Schaefer 1000 parcellation based on the 17-network Yeo RSN atlas^34^.

**Figure 3:**
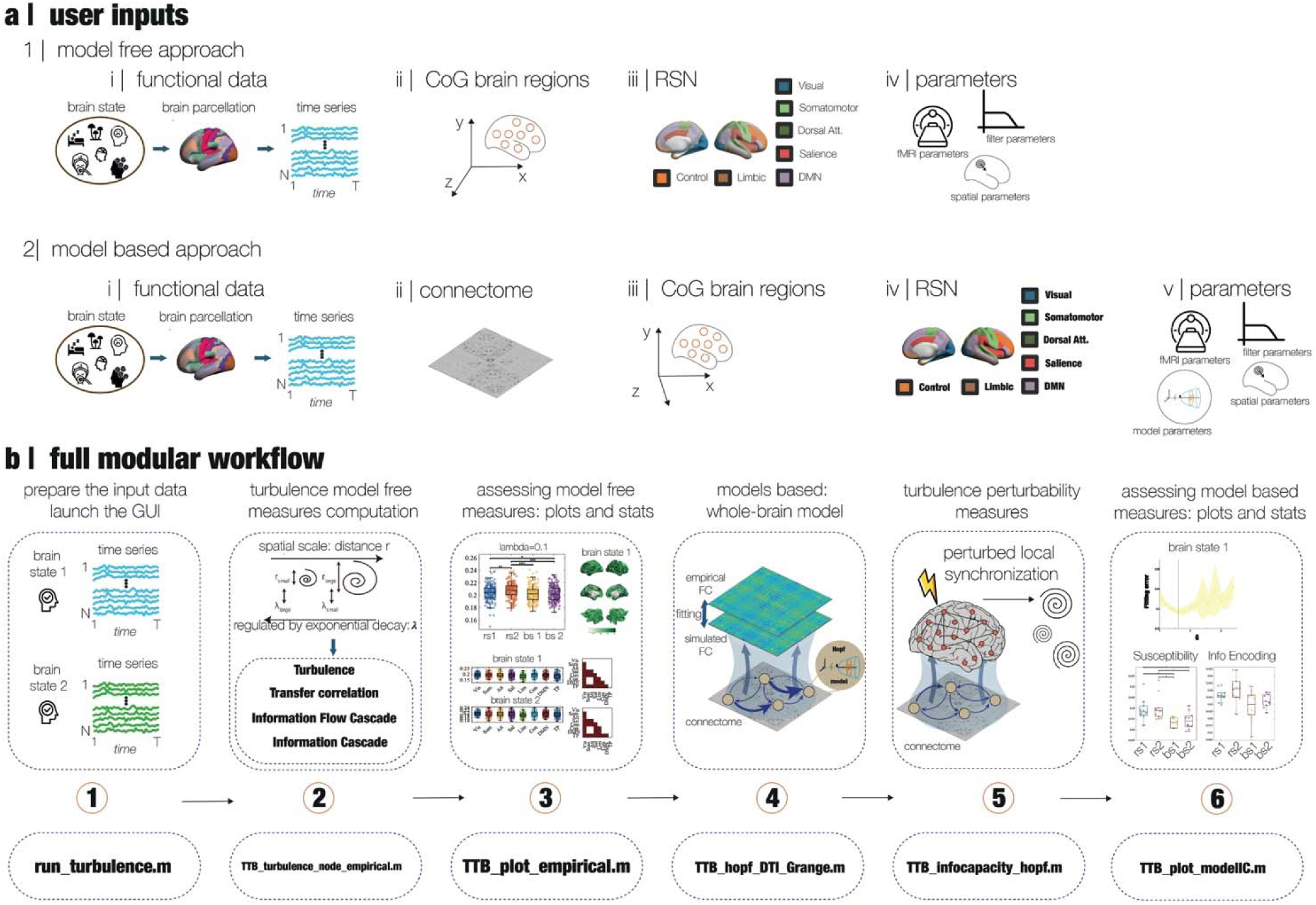
***The turbulent brain toolbox (TTB) workflow. a)*** *The general TTB workflow requires the following inputs to be provided by the user. For the model-free approach: i) functional MRI data for each participant and brain condition, mapped onto a high-resolution parcellation scheme (e.g., Schaefer 1000); ii) the spatial coordinates representing the centre of mass (CoG) for each brain parcel in the chosen parcellation; iii) the classification of each region within the parcellation according to its corresponding resting-state network (RSN), such as those defined by the Yeo 7-network atlas; iv) a collection of input parameters, categorized into fMRI-related (e.g., repetition time and number of time points), signal filtering settings, and spatial configuration parameters. For the model-based approach, the inputs are the same but adding a group-average structural connectome aligned to the same parcellation as the fMRI data and model parameters. **b)** In the full TTB workflow the input signal for each condition and participant is introduced to the model-free turbulence measures module. The model-free results are assessed and visualised in the next module. The whole-brain model for each brain condition and model perturbations for computing the turbulence perturbative measures (susceptibility and information encoding capabilities) are computed in the next two modules. Finally, the model-based results are quantified and plotted in the last module. Importantly, the TTB toolbox has a modular architecture composed of six modules, each one with a function supporting a specific step along the TTB pipeline (bottom row)*.

The complete TTB workflow including model-free and model-based approaches is shown in **Figure 3b**. In the first stage, fMRI data of each participant and each condition should be provided. fMRI data should have already undergone standard preprocessing pipeline, such as the Human Connectome Project (HCP) resting state minimal preprocessing described in detail on the HCP website. This pipeline includes standardized methods using FSL (FMRIB Software Library), FreeSurfer, and the Connectome Workbench software [30]. This standard preprocessing also includes correction for spatial and gradient distortions and head motion, intensity normalization and bias field removal, registration to the T1 weighted structural image, transformation to the 2-mm Montreal Neurological Institute space, and using the FIX artefact removal procedure [30,31]. These data should be upscaled to a fine-grained parcellation, which can include both cortical and subcortical regions. Importantly, the input should be a MATLAB cell array, with each column representing a brain condition and each row representing a participant. Each entry of the cell array should contain an **N×T** matrix, where **N** is the number of brain regions and **T** is the number of time points. In the second step, the complete set of model-free measures is computed for each participant in each condition: the turbulence at global level, RSN level and node-level and the transfer correlation for each indicated spatial scale (λ); the information flow cascade and information flow. In the third step, the toolbox computes the statistics for comparing model-free measures across brain conditions and delivers plots to visualize these results. In the fourth stage, a whole-brain model is created for capturing the brain dynamics of each brain condition included in the initial cell array provided by the user. Next step, the TTB computes the perturbative measures based on the variability of the local level of synchronisation (local KoP) when is perturbed for each brain condition. Finally, in the last stage the statistics and plots for the whole-brain model and turbulence perturbative measures are generated and displayed.

The TTB toolbox features a modular design comprising six modules, each containing functions that support a specific step in the full workflow presented in **Figure 3b**, which in turn can be used separately. The initial function is *run_turbulence.m,* which displays the main user-friendly GUI window where the users must provide all the inputs (see **Figure 3a**). The full set of turbulence model-free measures are computed by the *TTB_turbulence_node_empirical.m* function, which takes the input information and generates a *.mat* file with the results. The statistics and plots for the model-free results are performed by the *TTB_plot_empirical.m* function, which reads the results produced in the previous step and generates *.txt* output files summarizing the statistics of each comparison and *.fig* files with figures. Then, the whole-brain model is created by the *TTB_hopf_DTI_Grange.m* function obtaining an optimal coupling strength (G) value for each brain condition. Finally, the *TTB_Infocapacity_hopf.m* uses the output of the whole-brain model function and produces model perturbations and computes the susceptibility and information encoding capability measures. The results are plotted by *TTB_plot_modelIC.m* (**Figure 3b** **lower row)**.

### Tutorial

This section provides an overview of the multiple computations that the TTB toolbox supports (**Figure 4a**). The toolbox includes a graphical user interface (GUI) designed to guide users through the analysis pipeline via a structured, decision-based workflow. The interface follows a hierarchical logic tree that dynamically adapts to user selections, ensuring a streamlined and intuitive experience. Upon launching the toolbox by running the *run_turbulence.m* file, users are prompted to choose between a model-free and a model-based analysis approach. If the user selects the model-free approach, the GUI directly opens a data input window. This window allows users to specify the required inputs described in **Figure 3a**. Then the analysis will be focused on the modules 2 and 3, which are basically the turbulence empirical measures and the analysis of these results in terms of statistics and visualization plots. If the model-based approach is chosen, the GUI presents a second-level decision point, inquiring whether the model-free analysis has already been computed. If yes, the interface loads a dedicated window for inputting the model-based parameters, including the structural connectome and model configuration, applying modules 4 to 6 (see **Figure 3a**). If no, the user is asked whether fMRI data is available. Based on the response, the GUI presents the appropriate interface for either initiating the model-free step and the model-based approach fitted to the empirical data including the modules from 2 to 6 (this interface is shown in **Figure 4b**) or computing only model-based measures without fitting empirical data. This adaptive GUI structure minimizes user error and ensures that only relevant input fields are displayed at each stage. It was designed to be accessible to researchers and clinicians without requiring prior expertise in programming or computational modelling, while maintaining full analytical flexibility for advanced users.

**Figure 4:**
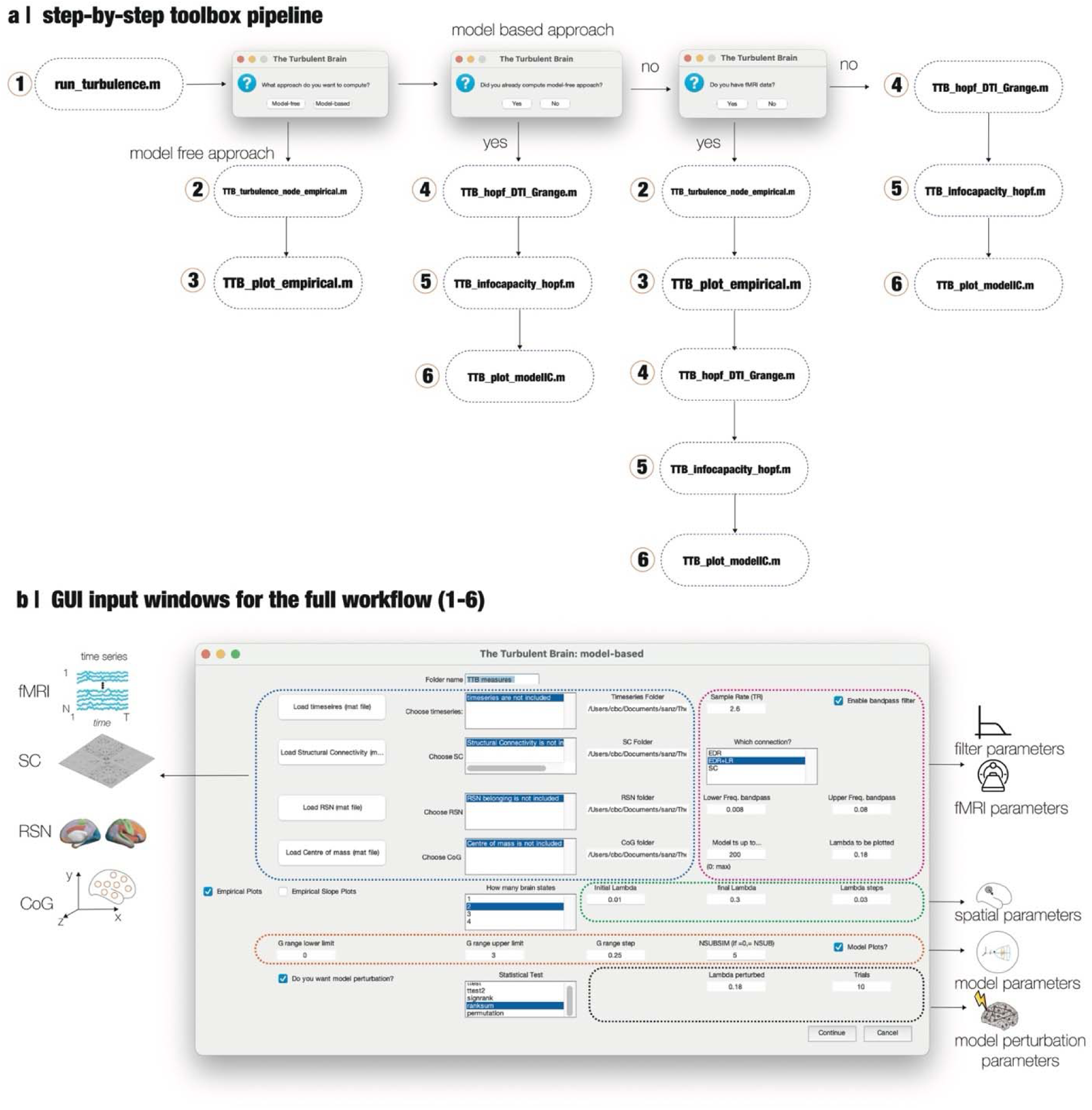
***The turbulent brain toolbox (TTB) step-by-step pipeline. a)*** Overview of the decision-based workflow guiding users through model-free and model-based analyses. The adaptive hierarchical structure ensures only relevant input fields are displayed at each stage. ***b)*** Full workflow interface indicated with five colour-coded panels: blue (empirical data inputs), red (configuration parameters), green (turbulence settings), orange (whole-brain model parameters), and black (perturbation parameters). This design streamlines analysis setup while remaining accessible to both novice and expert users.

**Figure 4b** illustrates the interface presented by the GUI when the user selects the full workflow, which includes both the model-free and model-based approaches. The input fields are organised into five main sections, visually delineated by coloured borders. The blue section are related with empirical data inputs including fields for loading the empirical data required for the analysis: fMRI data for each brain condition and participant; the group-average structural connectome (SC), the centre of mass (CoG) coordinates for each brain region in the selected parcellation; and the assignment of each brain region to one of the resting-state networks (RSNs). The GUI allows users to browse their local file system and select the corresponding *.mat* files from any directory. The red section is related to configuration parameters including the filtering range (lower and upper bounds) applied to the fMRI signals, the TR (repetition time) of the fMRI data, the number of time points to be used in the analysis and one spatial scale (λ) to make the empirical plots. The GUI allows for the selection of the connectivity scheme for constructing the whole-brain model, which includes: the Exponential Distance Rule (EDR), EDR plus long-range exceptions, and the actual empirical structural connectome (SC). The green section is related to turbulence parameter configuration, which requires the user to define the initial and final spatial scales (λ) as well as the step size for exploring this range. These values are used to compute the empirical turbulence measures across the selected spatial scales. The orange section, whole-brain model parameters, involves the configuration of the whole-brain model, specifically: the exploration range of the global coupling strength (G), defined by an initial value, a final value, and a step size and the number of simulations to run at each value of G. An option to indicate whether model-based plots should be generated. Finally, the black section related with perturbation parameters requires the spatial scale (λ) at which the perturbations will be applied and the number of repetitions. Then the GUI is also asking for defining a name for the folder where the analysis will be stored, the number of brain states to be included in the analysis, whether the user requires empirical plots and the statistical analysis to be performed in terms of paired,unpaired,parametric or non-parametric tests. Each section is designed to guide the user through the parameterization of their analysis pipeline, ensuring all necessary configurations are provided for the toolbox to function properly.

Once any of the alternatives is launched, the MATLAB command window will display the progress of the computations in terms of the number of subjects and brain states. Finally, when the computations are complete, the toolbox generates three types of files: i) .mat files containing the full set of results, ii) .fig files saving the figures generated with the toolbox, and iii) .txt files with the full statistical comparisons at the brain state level.

### Application case: movie watching

Here, we applied the full workflow of the TTB toolbox to a large-scale Human Connectome Project (HCP) neuroimaging data of the 176 individuals watching movies and resting (scanned with 7 T) for quantifying the information transmission flow changes in the brain. We leveraged this dataset that includes movie-watching Hollywood (h) and Creative Commons (cc) movies, and we explored the changes in brain during both contexts. In **Figure 5**, we gathered the results of the model-free analysis using the set of figures that the TTB generates. We computed the turbulence and information transfer for 9 spatial scales (from λ =0.03 to λ =0.3 in 0.03 steps) and displayed in **Figure 5** three of them λ =0.03, λ =0.15 and λ = 0.3 (for the complete set of scales see **Supp** Fig 1). We found that the turbulence increases at lower scales (higher λ) and decreased in high scales for both movie-watching, while for information transfer both movie watching conditions show a decrease comparing with the resting state (**Figure 5** **first row**). We then noticed that the Information Cascade significantly decreases between movie-watching and the resting state. Interestingly, these three measures showed significant differences between both movie-watching conditions, with less information transmission for the common creative movies. Finally, we display the node-level metastability and the turbulence at the functional network level for λ = 0.15 as an example. The node-level metastability is represented by brain renderings comparing the resting state and the common creative movie-watching condition, showing lower values of node-level metastability across the whole brain. The functional system-level turbulence analysis is performed across conditions as well as intra- and inter-condition comparisons. The statistics of these comparisons are represented by the lower diagonal matrices, with brown entries representing p < 0.05 (rank-sum test with false discovery rate (FDR) multiple comparison correction). Interestingly, the functional system analysis does not show clear differences across conditions, providing evidence that the differences between conditions could be related to a global brain information transmission effect.

**Figure 5:**
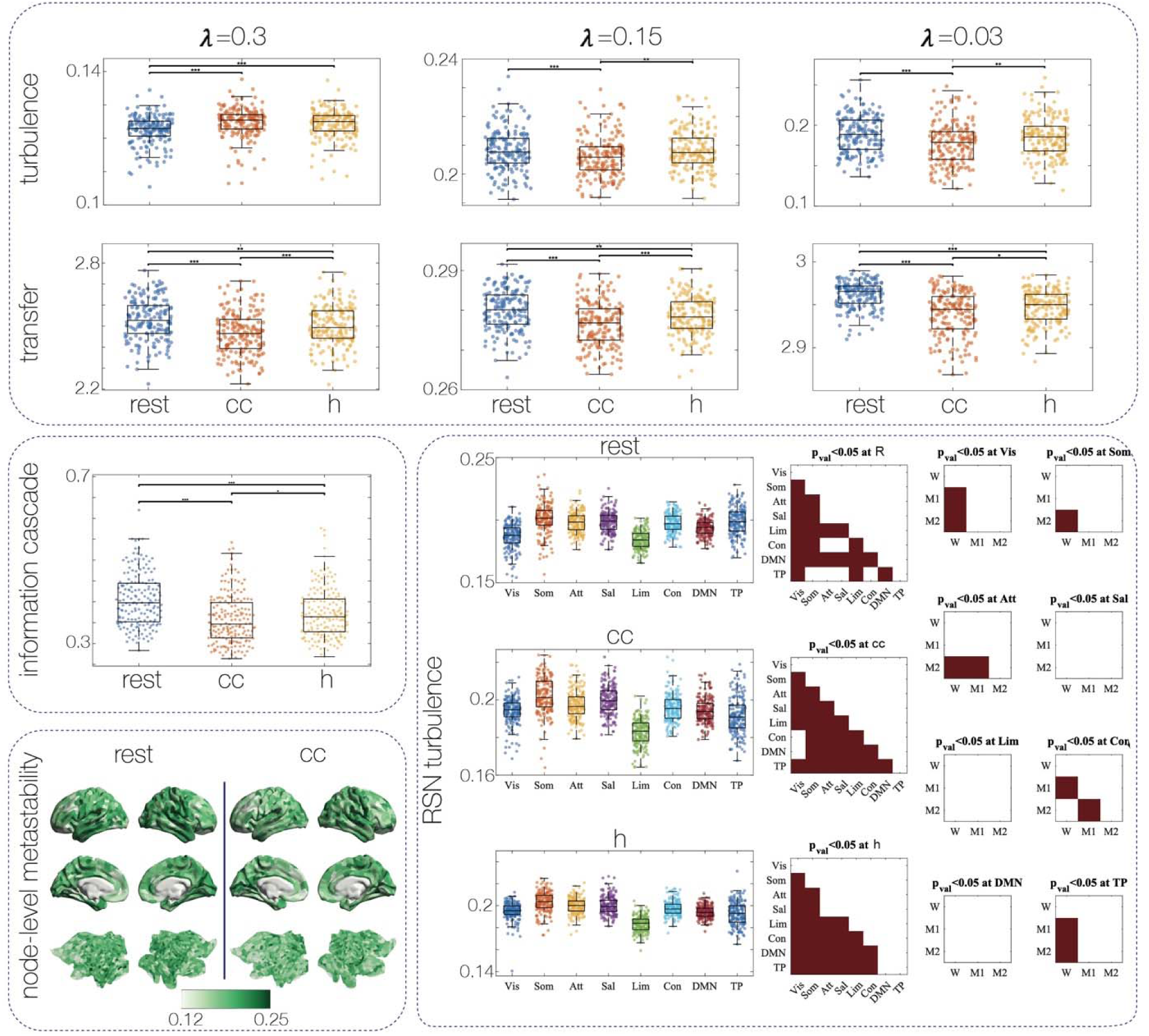
***The turbulent brain toolbox (TTB) application movie watching. Model-free results.*** *In the upper row the turbulence and information transfer comparison between resting-state and both movie-watching conditions groups (Comon Creative, cc, and Hollywood, h) for three spatial scales are displayed. In the middle row left, the Information cascade for the three conditions is shown.**Turbulence by RSN** - Functional system-level turbulence for spatial scale =0.15 (results showed a p-value threshold of <0.05). **Node level Metastability** - Node level metastability brain renderings for resting-state and Creative Commons (cc) conditions for spatial scale* λ *=0.15. We used is Wilcoxon rank sum test with false discovery rate correction for multiple comparisons. (Pval<0.001 with ***; 0.001<Pval<0.01 with **, and 0.01<Pval<0.05 with *)*.

After computing the model-free results we applied the model-based approach to the same data set. The first step is to build a whole-brain model for each condition. The TTB created a whole-brain model based on coupled non-linear oscillators and fitted to the group level empirical functional connectivity (FC) as a function of the distance (see Methods). The toolbox explores the user-indicated coupling strength parameter range (G) and computes the error fitting as the distance between the empirical and simulated FC for the number of simulations indicated by the user. **Figure 6** upper row displays the error fitting curve as a function of G for each condition and vertical lines indicates the G for the model working point, i.e. the minimum value of the fitting error, for each case. We found that the resting state condition shows the highest value of G (*G_rest_* = 2.0), then the Hollywood condition (*G_h_* = 1.6) and the lowest is for the common creative condition (cc) (*G_CC_* = 1.3). In the lower row of **Figure 6**, we present the results of the perturbative approach, quantifying the Information Encoding Capacity and Susceptibility for a spatial scale of (λ = 0.15). These measures were computed to assess the brain’s response to perturbations across the resting state, Hollywood (h), and Creative Commons (cc) movie-watching conditions. We observed significantly lower values for both Information Encoding Capacity and Susceptibility in the Creative Commons condition compared to the resting state (p < 0.001). Additionally, the cc condition showed significantly lower Information Encoding Capacity compared to the Hollywood condition (p < 0.001), but no significant difference was found for Susceptibility between the Creative Commons and Hollywood conditions. These findings indicate that the Creative Commons movie-watching condition is associated with reduced information encoding and perturbation responsiveness compared to the resting state, with specific differences in encoding capacity relative to the Hollywood condition.

**Figure 6:**
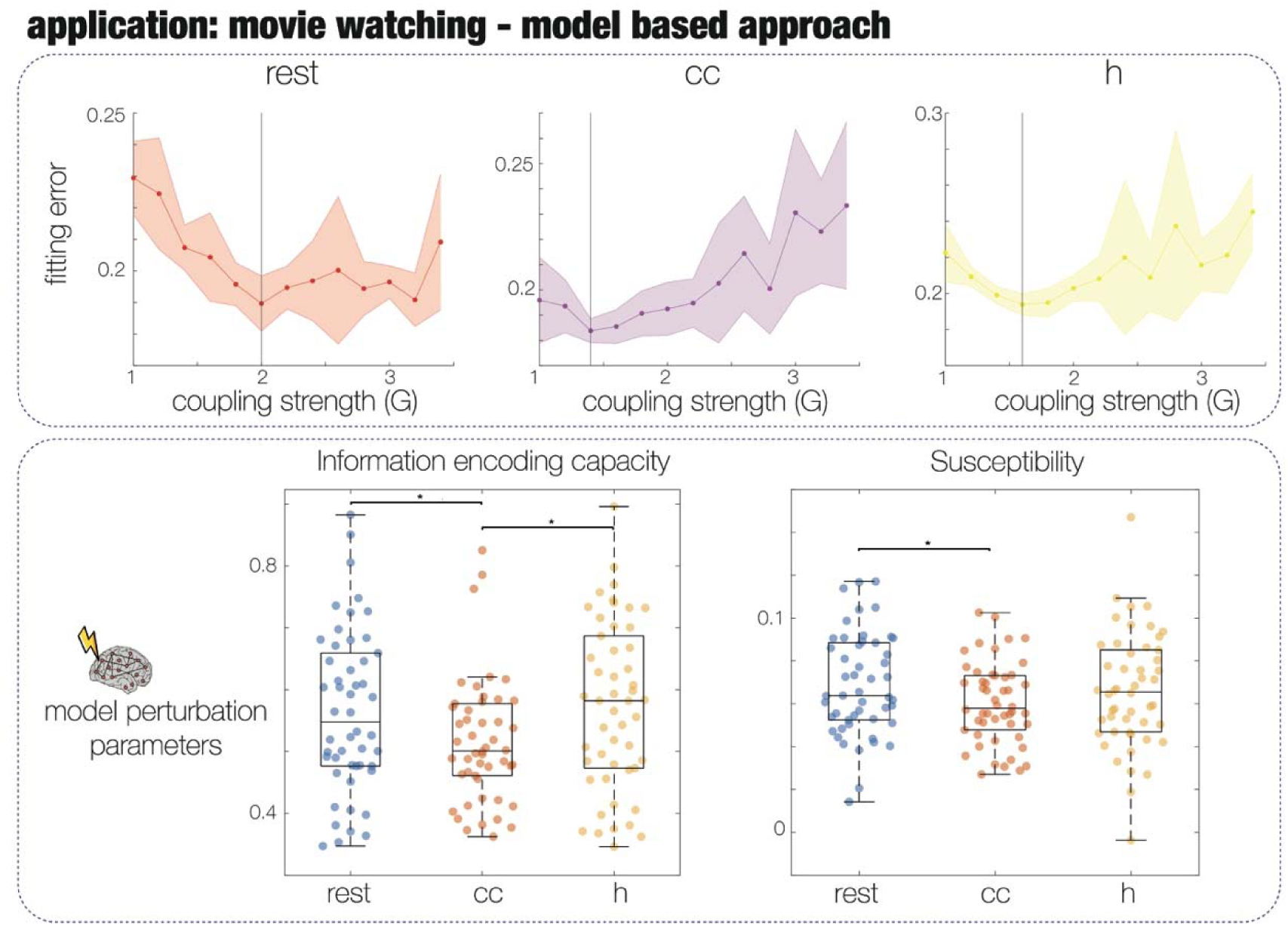
***The turbulent brain toolbox (TTB) application movie watching. Model-based results.*** *The upper row displays the whole-brain model error fitting value as a function of the coupling strength parameter (G) for each condition: resting state, common creative (cc) and Hollywood (h) movie-watching conditions. The vertical line in each plot indicates the G for the model working point, i.e. the minimum value of the fitting error. In the lower row presents the results of the perturbative approach in terms of both turbulence measures Information encoding capacity and Susceptibility for* λ *=0.15. We noticed that the cc condition presents significantly lower values for both measures compared with resting state and h conditions. The statistics we used is a Wilcoxon rank sum test with false discovery rate correction for multiple comparisons. (Pval<0.001 with ***; 0.001<Pval<0.01 with **, and 0.01<Pval<0.05 with *)*.

## Discussion

In this work, we introduced The Turbulent Brain Toolbox as an open-source MATLAB pipeline that enables both model-free and model-based analyses of turbulence in whole-brain dynamics. The toolbox provides an accessible framework for quantifying key turbulence-derived measures, including amplitude turbulence, information transfer, and information cascades, directly from empirical fMRI data, as well as for constructing and perturbing whole-brain Hopf models to estimate susceptibility and information encoding capacity. By lowering the technical barrier to applying these methods, TTB makes it possible for researchers and clinicians to explore how turbulent dynamics vary across brain states. As an illustrative application, we analysed ultra-high field fMRI data from the Human Connectome Project during resting-state and movie-watching^35^. Our findings showed that movie watching increased turbulence at fine scales while reducing information transfer and cascades relative to rest, with stronger reductions during Creative Commons compared to Hollywood films. Whole-brain modelling further revealed state-dependent shifts in coupling strength and lower perturbative capacity in the Creative Commons condition, highlighting that turbulence-based measures can sensitively differentiate brain states and capture the efficiency of large-scale information processing.

The turbulence framework, and the TTB as its implementation, are versatile across scales, modalities, and experimental contexts. At the macroscale, turbulence measures have been applied to fMRI data to distinguish global brain states such as sleep, disorders of consciousness^17^, traumatic brain injury^36^, and psychedelic states^18^, each showing characteristic shifts in the level of spatiotemporal variability. At finer temporal resolutions, the same formalism has been adapted to MEG^20^, capturing turbulence in fast oscillatory dynamics, while at the microscale, invasive recordings in hippocampal circuits have revealed signatures of turbulent cascades.^12^ Beyond modalities, turbulence has proven sensitive to both exogenous perturbations (e.g., pharmacological interventions^18^ or sensory stimulation such as movie watching here) and endogenous variations (e.g., meditation^17^ or menstrual cycle^37^). In particular, recent work showed how brain turbulence measures before pharmacological treatments in patients with major depressive disorder (MDD) is close related with the treatment outcome^38^. These diverse applications underline that turbulence is not restricted to a single type of data or experimental paradigm but instead provides a general language for quantifying how information is mixed and redistributed across the brain. As such, TTB serves as a unifying platform, allowing researchers to probe turbulence across conditions, datasets, and scales of organisation within a single framework.

Gathering these results together, we can identify that the turbulence framework is aligned with the growing focus on computational precision medicine as a promising avenue for uncovering the mechanistic principles governing both healthy and pathological brain dynamics^30–32^. Importantly, this framework not only propose the use of turbulence measures as fingerprints of individual brain states but also using the model-based branch of TTB as a route toward model-informed interventions. Perturbative analyses can reveal how different brain regions or networks contribute to maintaining or disrupting turbulent dynamics, thereby suggesting targets for neuromodulation strategies such as transcranial electrical stimulation (tES) or transcranial magnetic stimulation (TMS)^39,40^. While translation will require extensive validation, TTB establishes a practical framework to bridge empirical data, whole-brain modelling, and clinically relevant applications, paving the way for turbulence-based approaches in personalised medicine. This TTB capabilities is also supporting the move toward precision as a promising route in the field.

As a concrete application of TTB, we analysed fMRI data from the HCP project to investigate changes in the brain’s turbulent dynamics during movie watching compared to the resting state. Our findings from the model-free approach reveal distinct patterns in brain information transmission during movie-watching compared to the resting state. The observed increase in turbulence at lower spatial scales (higher λ) and decrease at higher scales (lower λ) during movie-watching suggests a scale-dependent modulation of brain dynamics, potentially reflecting heightened local processing demands during naturalistic stimuli. The consistent reduction in Information Transfer and Information Cascade across both Hollywood and Creative Commons movie-watching conditions compared to the resting state indicates a suppression of large-scale information flow, possibly due to focused attention on external stimuli. Notably, the Creative Commons movies exhibited lower information transmission, which may be attributed to differences in narrative complexity or emotional engagement compared to Hollywood movies. The lower node-level metastability during movie-watching, particularly for Creative Commons movies, suggests reduced dynamic flexibility across brain regions, potentially reflecting a more constrained functional state. However, the lack of clear differences in turbulence across conditions points to a global rather than network-specific effect, suggesting that movie-watching induces a broad reconfiguration of information transmission rather than localised network changes. Interestingly, this reduction of brain information transmission was observed during deep sleep and in disorders of consciousness patients^17^. At first glance, one might expect movie-watching to demand greater computational effort than resting, thereby engaging a steeper information transmission across the brain. Contrary to this intuition, our model-free analyses demonstrate the opposite effect. Previous work has observed that movie-watching is associated with a flattening of the brain’s hierarchical structure^41^, also related with brain states as deep sleep and disorders of consciousness^42,43^. This combined reduction in information transmission and hierarchical differentiation may help explain why film viewing is often experienced as a relaxing and restorative activity.

Furthermore, the model-based analysis using whole-brain models based on coupled non-linear oscillators provides additional insights into the underlying dynamics. The fitting of these models to empirical functional connectivity revealed distinct optimal coupling strengths (G) across conditions, with the resting state showing the highest coupling strength (*G_rest_* = 2.0), followed by Hollywood movie-watching (G_h_ = 1.6), and the lowest for Creative Commons movie-watching (*G_CC_* = 1.3). These differences suggest that the resting state supports stronger global synchronisation, potentially facilitating broader information integration, while movie-watching, particularly Creative Commons movies, may involve reduced coupling, reflecting a more fragmented or task-specific network configuration. Interestingly, this reduction of the coupling parameter strength was also found in disorders of consciousness patients^44^, traumatic brain injury^45^ and deep sleep^46^. The alignment of these model-based findings with the model-free results reinforces the notion that naturalistic stimuli modulate brain dynamics at both local and global scales, with varying degrees of synchronisation. These findings align with prior studies on naturalistic stimuli and highlight the utility of combining model-free and model-based approaches^41,47,48^, to capture the complex interplay of brain dynamics during naturalistic and resting conditions, offering a comprehensive view of how external stimuli shape global and local neural processing. Additionally, the perturbative approach revealed significantly lower Information Encoding Capacity and Susceptibility in the Creative Commons condition compared to the resting state, with only Information Encoding Capacity showing a significant reduction compared to the Hollywood condition. These findings suggest that the Creative Commons condition induces a more constrained neural state, particularly in terms of information encoding, potentially driven by lower external stimuli. The lack of significant differences in Susceptibility between Creative Commons and Hollywood conditions further indicates that perturbation responsiveness may be similarly affected by movie-watching, regardless of content type. Collectively, these findings highlight the utility of combining model-free, model-based, and perturbative approaches, as implemented in the TTB toolbox, to capture the complex interplay of brain dynamics during naturalistic and resting conditions, offering a comprehensive view of how external stimuli shape global and local neural processing.

Here turbulence is used in the context of neuroscience, but in a broader sense it describes a universal regime of nonlinear, high-dimensional systems far from equilibrium. Since turbulence emerges in a wide variety of natural and artificial systems - from weather and ocean currents, to plasmas, photonic devices, and even biological tissues, it has been conceived as a general principle for understanding how energy and information are transferred efficiently across scales. In neuroscience, this universality serves as a compelling lens to interpret brain activity, where turbulent cascades link local oscillations with global coordination. In this sense, the TTB is conceived as a bridge between turbulence theory and neuroscience, facilitating the translation of mathematical concepts from fluid dynamics into the analysis of large-scale brain activity. By integrating turbulence-based metrics with neuroscience and dynamical systems, the toolbox enables a mechanistic exploration of how turbulent dynamics underpin information processing in the brain.

Overall, this toolbox provides a workbench to connect brain turbulence with the wider landscape of complex dynamical systems, reinforcing both the biological relevance of turbulence and its role as a cross-disciplinary principle. At the same time TTB makes accessible for any user, without the requirement to have a background in computational modelling or mathematics, a set of computational tools for quantifying turbulence-related measures across neural signals and whole-brain models.

## Methods

### Ethics

The Washington University–University of Minnesota (WU-Minn HCP) Consortium obtained full informed consent from all participants, and research procedures and ethical guidelines were followed in accordance with Washington University Institutional Review Board (IRB) approval (Mapping the Human Connectome: Structure, Function, and Heritability; IRB #201204036).

### Participants

We used the data from the HCP using the 176 participants with data of movie-watching and rest (in 7 T), which were a subset of approximately 1200 in full HCP dataset. All participants (106 females and 70 males) were generally healthy young adults between 22 and 36 years old (mean age = 29.4 and SD = 3.3).

### Experimental protocol

Participants passively viewed a series of audiovisual movie clips in four functional 15 minutes runs consisting of four or five clips of varying length from around 1 to 4 minutes. In between each clip there was 20-s period of rest. The first and third runs contained clips from independent films (both fiction and documentary) made freely available under CC license on Vimeo, while the second and fourth runs contained clips from Hollywood films prepared by Cutt ^17^ing et al.^49,50^ The full description of both clips and the full experiment is provided in Finn et al ^35^. The movies were presented in a full-screen mode (size, 21.8° width × 15.7° height) and audio was delivered via Sensimetrics earbuds.

Four resting state scans (of approximately 16 min each) were acquired for the same participants. In the scans participants were instructed to keep their eyes open and maintain relaxed fixation on a bright cross-hair on a dark background in a darkened room. Given that the direction of phase encoding alternated between posterior to anterior and anterior to posterior across runs, we included only the runs anterior to posterior to provide a fair comparison between conditions.

### Neuroimaging data

A full detailed explanation of the participants acquisition protocol and data preprocessing in the HCP website (http://humanconnectome.org/). Below, we have briefly summarized this information.

### Structural data

Data were acquired using a customized 3-T Siemens Connectom Skyra scanner with a standard Siemens 32-channel radio frequency (RF)–receive head coil. For each participant, at least one three-dimensional (3D) T1w MPRAGE image and one 3D T2w SPACE image were collected at 0.7-mm isotropic resolution. In order to reconstruct a high-quality average structural connectivity (SC) matrix for constructing the whole-brain model (using the Schaefer1000 parcellation), we obtained multi-shell diffusion-weighted imaging data from 32 participants from the HCP database (scanned for approximately 89 minutes). The acquisition parameters are described in detail on the HCP website^51^. The connectivity was estimated using the method described by Horn and colleagues^52^. In summary, the data was processed using a generalized q-sampling imaging algorithm implemented in DSI studio (http://dsistudio.labsolver.org). A white-matter mask was produced from segmentation of the T2-weighted anatomical images, co-registered the images to the b0 image of the diffusion data using SPM12. For each of the 32 HCP participants, 200,000 fibres were sampled within the white-matter mask. Finally, the connectivity between the 1000 regions were averaged across participant to obtain a single high-quality structural connectivity included in the TTB as a source for building the model-based approach.

### Functional data

For each participant, HCP fMRI data were acquired using a 7-T Siemens Magnetom scanner with a Nova32 32-channel RF receive head coil, using the following parameters: 1.6-mm isotropic voxels, TR = 1000 ms, TE = 22.2 ms, flip angle = 45°, matrix = 130 × 130, FOV = 208 × 208 mm, 85 slices, multiband factor = 5, image acceleration factor = 2, partial Fourier sampling = 7/8, echo spacing = 0.64 ms, and bandwidth = 1924 Hz/Piels. The direction of phase encoding alternated between posterior to anterior and anterior to posterior across runs. For full details, see http://protocols.humanconnectome.org/HCP/7T/.

### Neuroimaging preprocessing for fMRI HCP

The preprocessing of the HCP resting state and task datasets is described in detail on the HCP website. The data are preprocessed using the HCP pipeline, which is using standardized methods using FSL (FMRIB Software Library), FreeSurfer, and the Connectome Workbench software^53,54^. This standard preprocessing included correction for spatial and gradient distortions and head motion, intensity normalization and bias field removal, registration to the T1 weighted structural image, transformation to the 2-mm Montreal Neurological Institute space, and using the FIX artefact removal procedure^54,55^. The head motion parameters were regressed out and structured artefacts were removed by ICA + FIX processing [independent component analysis followed by FMRIB’s ICA-based X-noiseifier ^56,57^]. Preprocessed time series of all grayordinates are in HCP Connectivity Informatics Technology Initiative (CIFTI) grayordinates standard space and available in the surface-based CIFTI file for each participants for resting state and each of the seven tasks.

We used a custom-made MATLAB script using the ft_read_cifti function [FieldTrip toolbox^58^] to extract the average time series of all the grayordinates in each region of the Schaefer 1000 parcellation^33^ with a total of 1000 cortical regions (500 regions per hemisphere), which are defined in the HCP CIFTI grayordinates standard space.

### Model-free

The mathematical formalism involved in the model-free measures that are included in the Toolbox are described below.

### Kuramoto Local order parameter

The time behaviour of the modulus of the Kuramoto local order parameter for a given brain area is named as the amplitude turbulence, *R_λ_*(*x̄,t*) and is mathematically defined as:

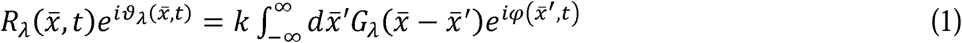

where *G_λ_* is the local weighting kernel 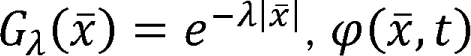 are the phases of the spatiotemporal data, *k* is the normalization factor 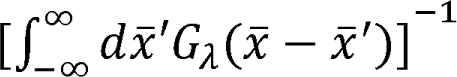 and λ is the spatial scaling.

Therefore, *R_λ_* defines local levels of 24ynchronization at a given scale, λ, as function of space, *x̄*, and time *t*. This measure captures what call the *brain vortex space*, *R_λ_*, over time, and can be related with the rotational vortices observed in fluid dynamics.

### Amplitude turbulence

The level of amplitude turbulence, *D*, is defined as the standard deviation across time and space of the modulus of local Kuramoto order parameter ®:

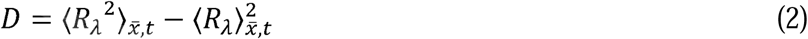

where the brackets 〈〉_*x̄,t*_ denotes averages across time and space.

### Information cascade flow and Information cascade

The information cascade flow indicates how the information travels from a given scale (λ) to a lower scale (λ − Δλ, where Δλ is a scale step) in consecutive time steps (*t* and *t* + Δ*t*). In this sense, the information cascade flow measures the information transfer across scales computed as the time correlation between the Kuramoto local order parameter in two consecutive scales and times:

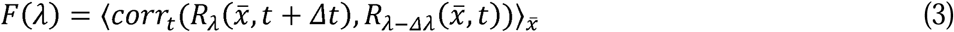

where the brackets 〈〉_*x̄,t*_ denotes averages across time and space. Then, the information cascade is obtained by averaging the information cascade flow across scales λ, which captures the whole behaviour of the information processing across scales.

### Information transfer

The spatial information transfer indicates how the information travels across space at a specific scale, λ. This measurement is computed as the slope of a linear fitting in the log-log scale of the time correlation between the Kuramoto local order parameter of two brain areas at the same scale as a function of its Euclidean distance ® within the inertial subrange:

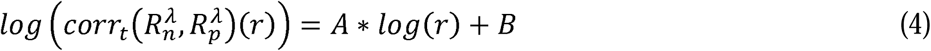

where *A* and *B* are the fitting parameters, and the first one, the negative slope, stands for the spatial information transfer.

### Node-level metastability

We computed the node variability of the local 25ynchronization as the standard deviation across time of the local Kuramoto order parameter as follows:

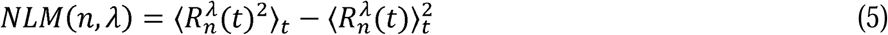

where the brackets 〈〉_*t*_ represent average values across time points.

Here, we used the discrete version of the node-level Kuramoto order parameter, with modulus *R* and phase ν, representing a spatial average of the complex phase factor of the local oscillators weighted by the coupling calculated through:

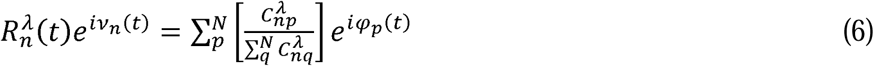

where *_p_*(*t*) are the phases of the spatiotemporal data, *N* is the total amount of nodes and *C^λ^_nq_* is the local weighting kernel between node *n* and *p*, and λ defines the spatial scaling:

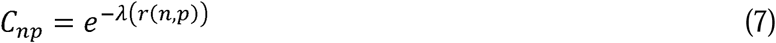

where *r*(*n,q*) is the Euclidean distance between the brain areas *n* and *p* in MNI space.

To compare the node-level metastability statistics, we collected the 1000 nodes values for all participants in each group and generated the distributions. Then, we compared across states the distributions using the Kolmogorov-Smirnov distance between them. The Kolmogorov–Smirnov distance quantifies the maximal difference between the cumulative distribution functions of the two samples, where larger values stand for more significant differences between both distributions.

#### 2.1.2 Model-based

We constructed whole-brain dynamical models based on the normal form of a supercritical Hopf bifurcation (also known as Stuart-Landau)^26^. This type of bifurcation can change the qualitative nature of the solutions from a limit cycle that yields self-sustained oscillations towards a stable fixed point in phase space. This whole-brain computational model is characterised by a series of model parameters that rules the global dynamical behaviour. One of them is the multiplicative factor, *G*, representing the global conductivity of the fibres scaling the structural connectivity between brain areas, which is assumed to be equal across the brain^26,59^. The other relevant parameters are the local bifurcation parameter (*a_j_*), which rules the dynamical behaviour of each area between noise-induced (*a* < 0), self-sustained oscillations (*a* > 0) or a critical behaviour between both (*a* ∼ 0). The bifurcation parameter is fixed critical regime (*a* =-0.02), where has been demonstrated the model present optimal behaviour^28^ ^29^. The model parameter G is 26ptimized to better fit the empirical functional connectivity as a function of the distance, *r*, within the inertial subrange. The models consisted of J brain areas from a given brain atlas. The underlying anatomical matrix *C_np_* was added to link the brain structure and functional dynamics and was obtained by measuring the exponential distance rule as defined in Equation (7). The local dynamics of each brain area is mathematically described as:

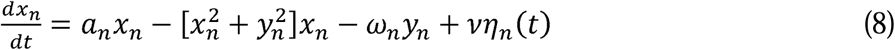

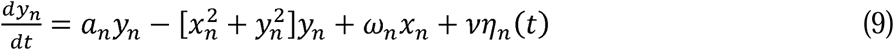

where η*_n_*(*t*) is additive Gaussian noise with standard deviation + D 0.01, *a_n_* is the bifurcation parameter of each node (we considered equally for all brain regions in this Toolbox), which allows the system to oscillate with frequency *f_n_* = ω*_n_*/2π. The frequency ω*_n_* of each brain area was estimated from the empirical fMRI data as the peak of the power spectrum.

Finally, the whole-brain dynamics was defined by the following set of coupled equations:

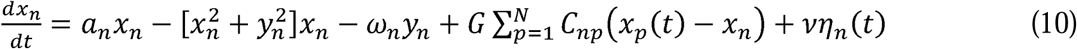

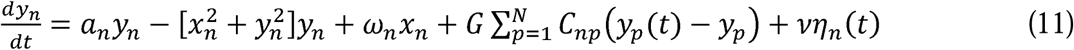

Where the global coupling factor G, scaled equally for each brain area, represents the input received in region *n* from every other region *p*.

### Functional Connectivity Fitting

An adaptation of Kolmogorov’s structure-function is proposed, and the variable *u* was applied to the BOLD signal of the data, instead to the velocity field of the fluid. Thus, the functional correlations between each pair of brain areas with equal Euclidean distance and can be defined as:

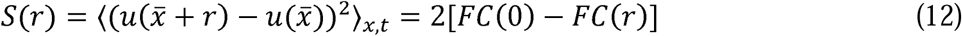

where *FC* is the spatial correlations of two points separated by a Euclidean distance r, which is given by:

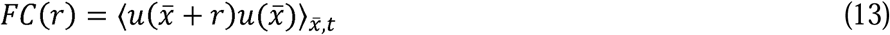

where the symbol 〈〉_*x̄,t*_ refers to the average across the spatial location x of the brain areas and time. Thus, the structure functions characterise the evolution of the functional connectivity (FC) as a function of the Euclidean distance between nodes at the same distance, which is different from the usual definition of FC that does include distance. The fitting between the empirical and simulated FC is defined as the Euclidean distance between both matrices within the inertial range as defined in Deco et al^9^.

### Susceptibility

How the brain reacts to external stimulations is quantified by the susceptibility measure of the whole-brain model following the definition proposed by Daido^60^ and implemented in previous works^17,38^. For each coupling strength value, G, the Hopf model was perturbed by randomly changing the local bifurcation parameter, *a_n_*, in the range [-0.02: 0]. The sensitivity of the perturbations on the spatiotemporal dynamics was calculated by measuring the changes in modulus of the local Kuramoto order parameter as:

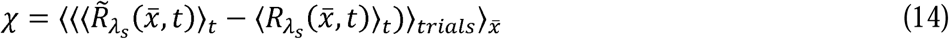

where *R̃_λ_s__*(*x̄,t*) corresponds to the perturbed case, the *R̃_λ_s__*(*x̄,t*) to the unperturbed case, and 〈〉_*t*_ and 〈〉_*trials*_ and 〈〉_*x̄*_ to the average across time, trials, and space, respectively.

### Information encoding capability

The information encoding capability captures how the external stimulations are encoded in whole-brain dynamics. The information capability, I, was defined as the standard deviation across trials of the difference between the perturbed *R̃_λ_s__*(*x̄,t*) and unperturbed *R_λ_s__*(*x̄,t*) mean of the modulus of the local Kuramoto order parameter across time *t*, averaged across all brain areas *n* as:

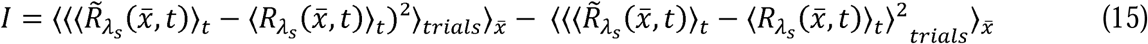

where the brackets 〈〉_*t*_, 〈〉_*trials*_ and 〈〉_*x̄*_ denote the averages defined as above.

## Acknowledgment

Y.S.P. was supported by the project NEurological MEchanismS of Injury, and Sleep-like cellular dynamics (NEMESIS) (ref. 101071900) funded by the EU ERC Synergy Horizon Europe. JV is supported by the project NEurological MEchanismS of Injury, and Sleep-like cellular dynamics (NEMESIS) (ref. 101071900) funded by the EU ERC Synergy Horizon Europe. A.E. was supported by the European Union’s Horizon Europe research and innovation programme under the Marie Skłodowska-Curie Actions (ID: 101207460, NEUROCONTRA, HORIZON-MSCA-2024-PF-01-01), by the project eBRAIN-Health - Actionable Multilevel Health Data (ID: 101058516) funded by the EU Horizon Europe, and by the project NEUROFEM funded by La Fundació la Marató de TV3 (ID: 202410-30-31). M.L.K. is supported by the Centre for Eudaimonia and Human Flourishing (funded by the Pettit and Carlsberg Foundations) and Center for Music in the Brain (funded by the Danish National Research Foundation, DNRF117). The funders had no role in study design, data collection and analysis, decision to publish or preparation of the manuscript. G.D. is supported by Grant PID2022-136216NB-I00 funded by MICIU/AEI/10.13039/501100011033 and by “ERDF A way of making Europe”, ERDF, EU, Project NEurological MEchanismS of Injury, and Sleep-like cellular dynamics (NEMESIS) (ref. 101071900) funded by the EU ERC Synergy Horizon Europe, and AGAUR research support grant (ref. 2021 SGR 00917) funded by the Department of Research and Universities of the Generalitat of Catalunya.

## Notes

### Competing Interest Statement

The authors have declared no competing interest.

